# An updated DNA barcoding tool for *Aloe vera* and related CITES-regulated species

**DOI:** 10.1101/2024.07.09.602761

**Authors:** Yannick Woudstra, Paul Rees, Solofo E. Rakotoarisoa, Nina Rønsted, Caroline Howard, Olwen M. Grace

**Author notes:** Correspondence: Yannick Woudstra.

## Abstract

DNA barcoding has revolutionised the identification of illegally traded material of endangered species as it overcomes the lack of resolution encountered with morphological identification. Nonetheless, in recently evolved and highly diverse clades, such as the relatives of *Aloe vera*, the lack of interspecific sequence variation in standardised markers compromises the barcoding efficacy. We present a new DNA barcoding tool using 189 nuclear markers, optimised for aloes (Asphodelaceae, Alooideae). We built a comprehensive sequence reference dataset from taxonomically verified sources for >300 species and validated its reliability for identification using phylogenomic inference. Seven anonymised samples from verified botanical collections and ten plants seized at London Heathrow Airport were correctly identified to species level, including a critically endangered species from Madagascar. Commercially purchased samples were confirmed to be the species as advertised. An accurate, reliable DNA barcoding method for aloe identification introduces new assurance to regulatory processes for endangered plants in trade.

## Introduction

International wildlife trafficking is having a severely negative impact on the survival of endangered species and is one of the main drivers of biodiversity loss^1^. This illegal market is valued at up to $216 billion^2^ and includes plant groups such as rosewoods^3^ (several tropical hardwood genera), cycads^4^ (Cycadaceae, Zamiaceae), and succulents. National legislation and task forces (https://unitedforwildlife.org/), along with international legislation such as the Convention on International Trade of Endangered Species (CITES; https://cites.org/), provide a legal framework for regulating international trade in targeted species by restricting transport across international borders. Proper implementation necessitates accurate and robust species identification of transported samples, yet traded material is mostly sterile and often processed, hence lacking diagnostic morphological characters ordinarily used for species identification.

DNA sequencing technologies have helped to overcome these problems through DNA barcodes^5^. These standardised genomic regions can be used to determine the taxonomic identity of a sample when compared to a curated reference dataset^5,6^ (e.g., the Barcoding of Life Data System (BOLD), http://www.barcodinglife.org). For animals, the cytochrome oxidase 1 gene does this effectively^7^ and for plants a combination of plastid (*matK, rbcL*) and nuclear ribosomal (ITS) markers were proposed as the standard^6^. These markers were successfully used in monitoring the illegal trade of timber products such as mahogany (Meliaceae) and laurel (Lauracaeae)^8^. However, insufficient sequence variation in recently diverged and diverse clades such as rosewoods^3^ (±300 species) and aloes^9^ (±600 species) require alternative methods. Full plastid genome sequences, obtained through genome skimming^10^ have improved identification in several plant groups (e.g. *Araucaria*^11^ and *Rhododendron*^12^), but do not resolve species boundaries in others^6^. More variable DNA sequences from the nuclear genome^13^, especially low-copy nuclear genes^14^, are now obtainable independent of genome size^15^ and DNA degradation level^16^, making them ideally suited for identifying diverse clades and traded plant material^17^.

Diagnostic tools are urgently needed for the succulent-leaved aloes (Asphodelaceae, Alooideae), which are threatened with extinction despite being cultivated for the horticultural and cosmetic industries^18^. This is particularly worrying in Madagascar, a centre of unique diversity in aloes^19^, where more than half of the endemic species are (critically) endangered (Figure 1). International trade in aloes is regulated by CITES legislation^20^ since illegal and unsustainable wild harvesting could be catastrophic for the survival of several species, but species identification is problematic. Full plastid genome sequences were successfully used to resolve species relationships in the small tree aloe genus *Aloidendron*^21^ (6 species) but have shown less promise in *Aloe* (±600 species). With the development of a target capture tool for aloes^15^, low-copy nuclear genes can now be sequenced cost-effectively from both fresh and more degraded (processed) material.

**FIGURE 1:**
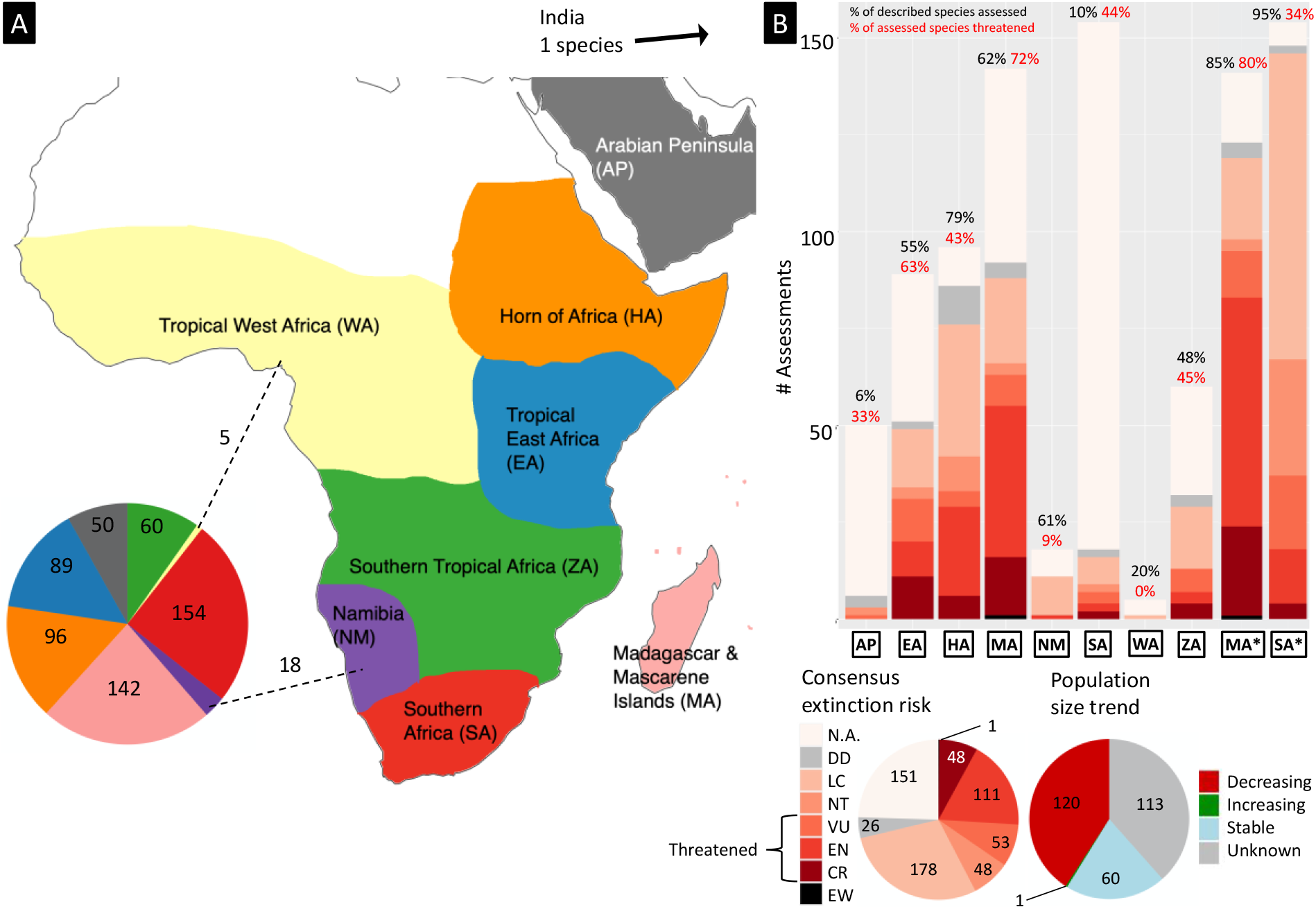
Extinction risk overview of the aloes (Asphodelaceae – subfamily Alooideae – genus *Aloe* L. and related genera; 616 species^22^). **A** – Main geographic areas of distribution for aloes: AP – Arabian Peninsula; EA – Tropical East Africa; HA – Horn of Africa; MA – Madagascar & Mascarene Islands; NM – Namibia; SA – South Africa; WA – Tropical West Africa; ZA – Zambezia or Southern Tropical Africa. Pie chart corresponds to the number of described species found in each region, based on the most recent species descriptions dataset (The World Checklist of Vascular Plants^x^, https://powo.science.kew.org). **B** – The distribution of extinction risk for aloes in each geographic region, based on the latest update of the IUCN Red List (2023-1^23^, https://www.iucnredlist.org/); separate distributions for Madagascar and South Africa are plotted where IUCN Red List data was complemented with pending assessments for Madagascar (S. Rakotoarisoa, Kew Madagascar Conservation Center) and published SANBI Red List assessments for South Africa^24^ (South African National Biodiversity Institute, http://redlist.sanbi.org/). Percentages above the bars indicate the proportion of species in each region that has been assessed (black), where Data Deficient (DD) assessments are not counted, and the proportion of assessed species that are threatened, the sum of assessments in categories VU, EN, and CR. Combining these datasets, a consensus extinction risk is visualised in a pie chart cover all 616 species of aloes. For officially published IUCN Red List assessments, the distribution of population size trend is also visualised in a pie chart. Extinction risk assessments are according to the IUCN Red List Categories and Criteria (IUCN Species Survival Commission (SSC) 2012): DD – Data Deficient; LC – Least Concern; NT – Near Threatened; VU – Vulnerable; EN – Endangered; CR – Critically Endangered; EW – Extinct in the Wild.

Here we present a DNA barcoding tool for the aloes, to aid in the international conservation of these threatened succulents. Through phylogenomic analysis with target capture^15^ we built an extensive reference dataset of 189 low-copy nuclear genes in 293 *Aloe* taxa and at least one member of every other genus in the Alooideae subfamily. The discriminatory power of the tool was validated with anonymised curated samples and was further tested on store-bought *Aloe* leaves and plants confiscated at the United Kingdom border (London Heathrow Airport). Our tool reliably identifies important conservation target species, such as critically endangered aloes from Madagascar, and establishes reliable and accurate DNA barcoding in this economically important plant group.

## Methods

The following is a summary of the methodologies used for this study. A full detailed description can be found in the Supporting Information.

### 1. Reference dataset

A comprehensive reference sequence dataset of aloes (Asphodelaceae, Alooideae) was compiled using taxonomically verified samples from botanical collections at the Royal Botanic Gardens, Kew (RBG Kew) and donations from a variety of natural history institutions (online supporting material). Both historical (herbarium) and living plant collections were consulted. We ensured that all currently recognised taxonomic and phylogenetic clades of the genus *Aloe* were included^19^, as well as smaller genera that make up the Alooideae subfamily^25^. The sampling comprised 375 species of *Aloe*, four species of *Aloidendon*, three species of *Aloiampelos*, and one species each of *Aloestrela, Aristaloe, Astroloba, Haworthia, Haworthiopsis, Gonialoe, Kumara* and *Tulista*. We sampled *Bulbine frutescens* (subfamily Asphodeloideae) as an outgroup. Twenty-eight samples were obtained during the design process for our Alooideae target capture tool^15^.

DNA was extracted from silica-dried leaf material from living plant collections and from historical leaf and/or flower material from herbaria. High molecular weight DNA was sheared to the appropriate fragment size using ultrasonication. Short-read sequencing libraries were created with dual index barcodes, combined in equimolar pools based on fragment size distribution, enriched using the Alooideae target capture tool^15^ and sequenced on Illumina® HiSeq X lanes.

Using a customised bioinformatic pipeline (Supporting Information for details), raw sequencing data were quality-filtered, assembled into nuclear target loci and aligned in locus-specific multiple-sequence-alignments (MSAs), which were cleaned for phylogenetic purposes. To provide sequence quality assurance and consistency, only samples with ≥50% total target recovery were included in the reference dataset. Potentially paralogous loci (genes comprising multiple highly similar copies from the same haploid genome) were removed from the dataset. Phylogenomic tree topologies were estimated from the MSAs with two methodologies: a maximum likelihood species tree from a concatenated supermatrix and a coalescent-based summary species tree from individual maximum likelihood gene trees.

### 2. Barcoding tool validation

To test the sensitivity of the Alooideae target capture barcoding tool, we enriched samples of unknown or uncertain identity. One of us (PR) prepared ten anonymised samples from previously unsampled accessions from the living collections at RBG Kew. Two phylogenomic inferences (see above) were contrasted with a more traditional method using pairwise genetic distances for sample identification. We combined the nuclear target sequences of the anonymised samples with the reference dataset and reran the custom pipeline to generate new MSAs. Phylogenomic inferences were then repeated for the updated dataset and sample identities were determined as the taxon with which the unknown sample is recovered in a monophyletic clade, with the node support value indicating the confidence in this identity. In the case of non-monophyly, the closest relative of the unknown sample was noted, if applicable. Sequences of these newly identified samples were verified by PR then added to the reference dataset.

### 3. Identification of unknown samples

The validated tool was then used to identify ten *Aloe* plants, identified to genus rank only (Figure 2), seized by border officials at London Heathrow Airport and quarantined at RBG Kew, the UK CITES Flora authority. Additionally, we obtained three fresh leaf samples from the UK domestic and international market to test the tool on traded samples. Two samples were purchased as *Aloe vera* leaves from organic supermarkets in London, United Kingdom, and one sample was purchased online as *Aloe arborescens* Finally, we applied the tool to the curation of unidentified specimens at RBG Kew using six plants in the living collections and ten undetermined wild-collected herbarium specimens, all of which were identified to genus rank only. Barcoding gaps, defined as the difference between inter- and intra-specific genetic distances^26^, were determined to further assess the confidence of species identifications.

**FIGURE 2:**
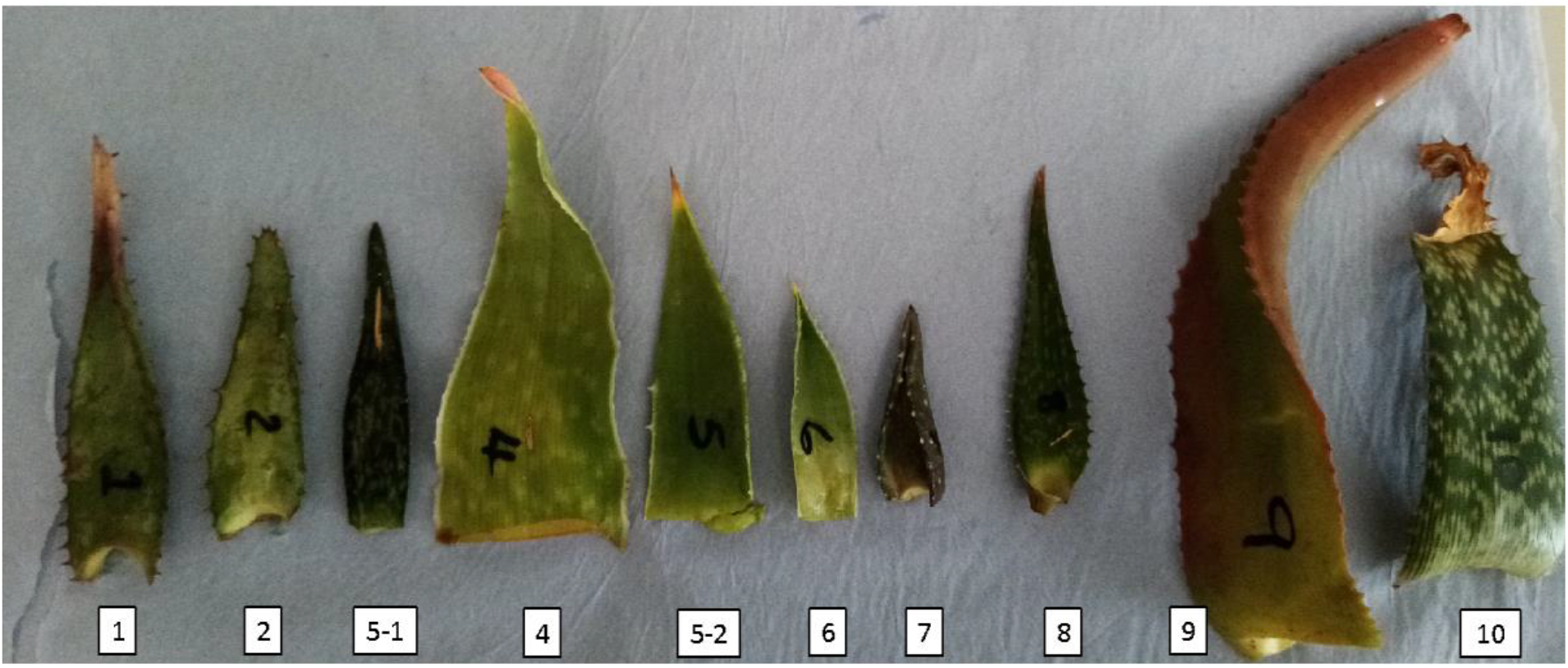
Leaf samples of ten aloe plants seized by border control at London Heathrow Airport, identified to species rank (“CITES” samples in Table 1) using phylogenomic placement of target capture DNA sequences, for which importation into the United Kingdom was deemed in conflict with international CITES restrictions (e.g., material not belonging to *Aloe vera* or *Aloe ferox*).

### 4. Conservation target assessment

Identifying the *Aloe* species among CITES interceptions has the potential to reveal real-time trends in illicit transport of species and refine conservation targets. We mapped available extinction risk assessments for aloes from the IUCN Red List^23^ (https://www.iucnredlist.org/), complemented by new (pending) assessments for Madagascan taxa (by SR), and SANBI Red List assessments^24^ for South African taxa (http://redlist.sanbi.org/) to the phylogenomic framework obtained with our reference sequence dataset. Species identified among our CITES-restricted samples were then manually highlighted on the tree to visualise phylogenomic patterns in extinction risk.

**TABLE 1:**
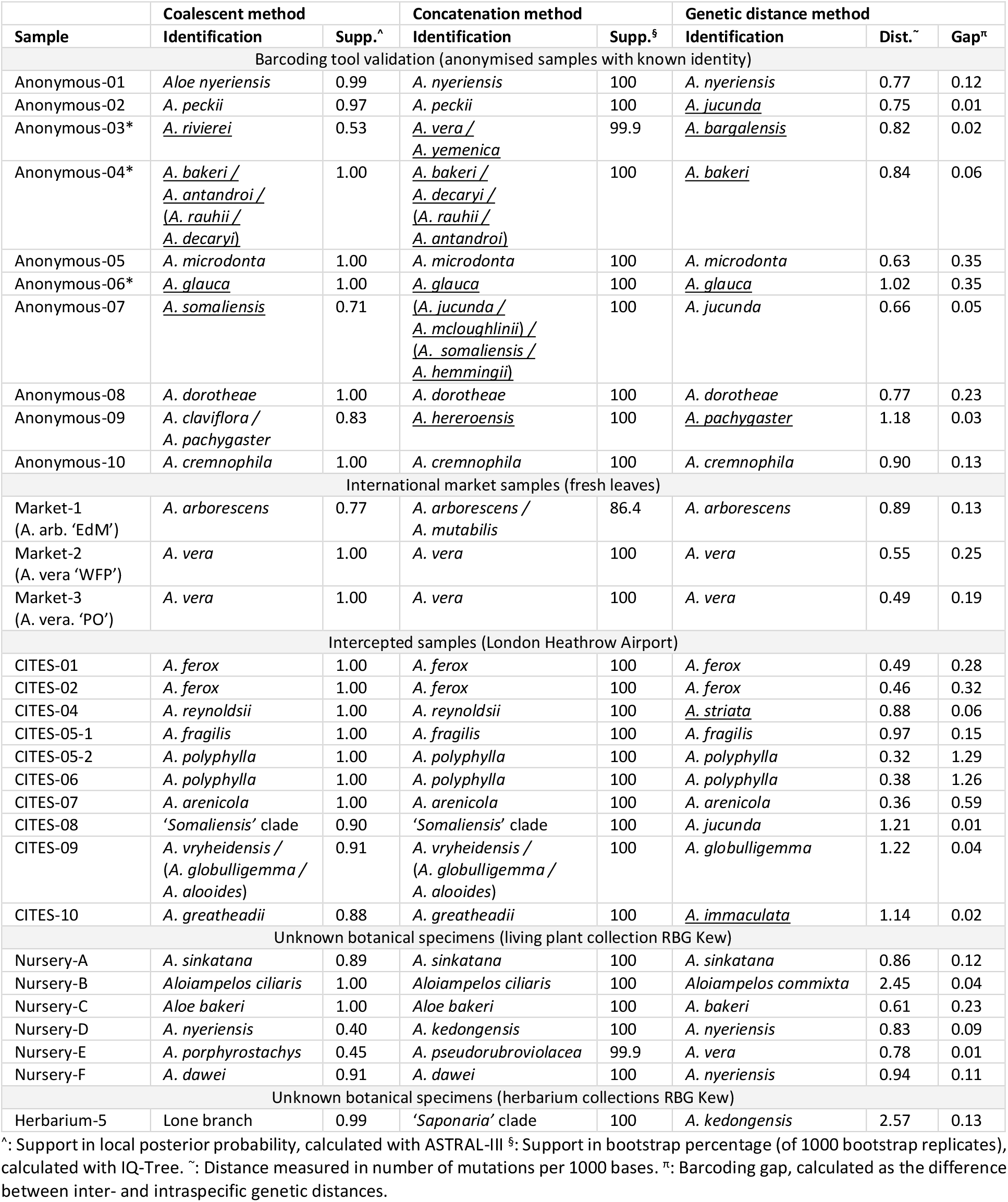
Details of the DNA barcoding results, following the order of our workflow (Figure 2): starting with the validation results (“Anonymous-”), followed by the international market (“Market-”) and trade (“CITES-”) samples, and ending with the unidentified specimens from living (“Nursery-”) and preserved (“Herbarium-”) botanical collections. Misidentifications are underlined, with * indicating the absence of reference material for the true species. The presence of multiple species names in an identification indicates a species complex and the denotation follows the phylogenetic relationships within the sister clade to the unknown sample.

## Results

The following is a summary of the results of this study. A detailed overview of sequencing results and DNA barcoding inferences are available in the Supporting Information.

### 1. Reference dataset

The reference sequence dataset for Alooideae comprised 310 samples representing 285 species of *Aloe* (including five subspecies and four varieties), four species of *Aloidendron*, three species of *Aloiampelos* and one species each of the remaining Alooideae genera, as well as outgroup *Bulbine frutescens*. The full dataset was trimmed from 189 to 173 target loci, after removing paralog-containing alignments and poorly recovered loci (<200 samples represented).

The tree topologies obtained with the concatenation and coalescent methods both showed the same relationships between Alooideae genera. Most of the *Aloe* topology was consistent in both estimation methods, with large monophyletic clades following strong geographical patterns (e.g., several South African clades, Madagascar, Namibia, Zambezia, Tropical East Africa, and the Horn of Africa / Arabian Peninsula; Figure 3). There were, however, considerable differences between the two topologies in the placement of taxa sharing the MRCA with all *Aloe* species, indicating incomplete lineage sorting.

**FIGURE 3:**
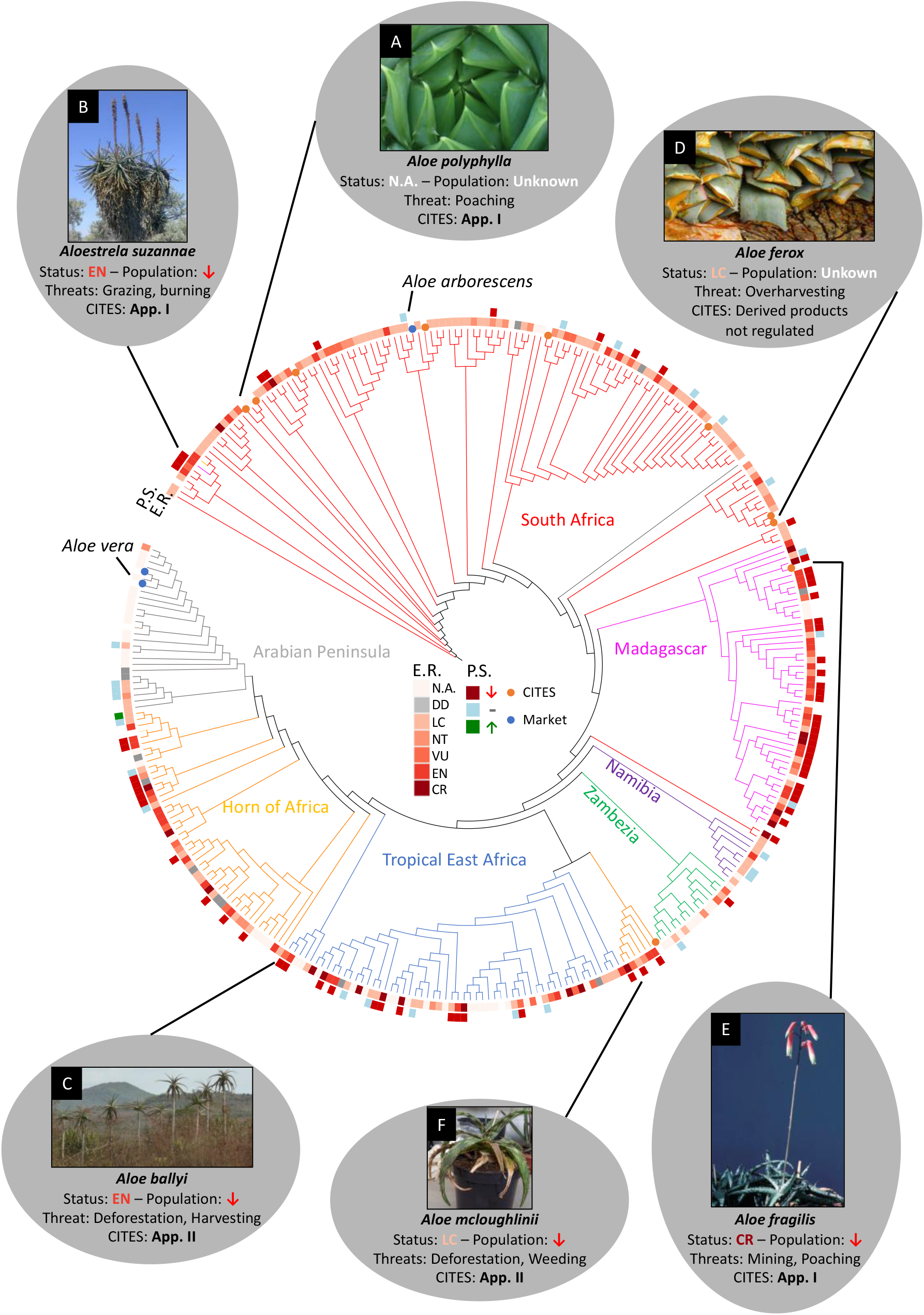
*Aloe* customised DNA barcoding in a conservation context. Phylogenomic framework of Alooideae, based on the updated reference dataset, with identifications of intercepted illegal trade material (orange dots) and international market samples (blue dots). Extinction risk (“E.R.”) for each species is plotted according to our consensus dataset, comprising the most recent (2023-1)^23^ IUCN Red List assessments, complemented with new assessments for Madagascar and SANBI Red List assessments for South Africa (Figure 1A). The population size trend (“P.S.”) is indicated as decreasing, stable, or increasing. Phylogenetic clades are coloured according to the predominant geographic distribution area (Figure 1B for definitions). Six examples of aloes with economic importance and/or dire conservation risk are highlighted. Image credits: A & F – Y. Woudstra; B – S. Rakotoarisoa; C – P. Pavelka, Royal Botanic Gardens, Kew; D – T. Hoffman, taken from Melin et al. (2017)^27^; E – U. Eggli, taken from Newton (2020)^19^.

### 2. Barcoding tool validation

The sensitivity of the Alooideae target capture barcoding tool in identifying *Aloe* DNA extracts to species rank was confirmed by the fully accurate taxonomic clade placement and 60% accurate species placement of anonymised curated samples (“anonymous-” in Table 1) using the coalescent phylogenomic method. Of the four samples where species placement was ambiguous, three (Anonymous-03, -04 and -06) belonged to species that were not yet present in our reference dataset (*A. pendens, A. millotii* and *A. lineata*, respectively) and were henceforth added to update the dataset. The remaining sample (Anonymous-07) belonged to the *Aloe somaliensis* species complex. A lower success rate of 50% species rank identification was observed using the phylogenomic concatenation method and the genetic distance method. We therefore considered the phylogenomic coalescent version of our DNA barcoding tool validated on the basis that every sample was assigned to the correct clade, with gene tree conflict and phylogenetic support giving clear and reliable indications of caution.

### 3. Identification of unknown samples

A total of twenty unknown samples were identified with our DNA barcoding tool for aloes, of which seventeen were confidently identified to a single species. Identifications of intercepted CITES-regulated samples were highly supported and fully congruent between the two phylogenomic methodologies applied (Table 1). The accuracy appears to be reduced only in assigning species-level identification to taxa in species complexes, such as that of *Aloe somaliensis* (sample CITES-08). Barcoding gaps are generally low in species complexes (Table 1), reflecting low genetic distances between species in the clade, but are well established in the economically important species (Figure 4). The three leaf samples bought on the international market were all determined to be the species indicated by the sellers.

**FIGURE 4:**
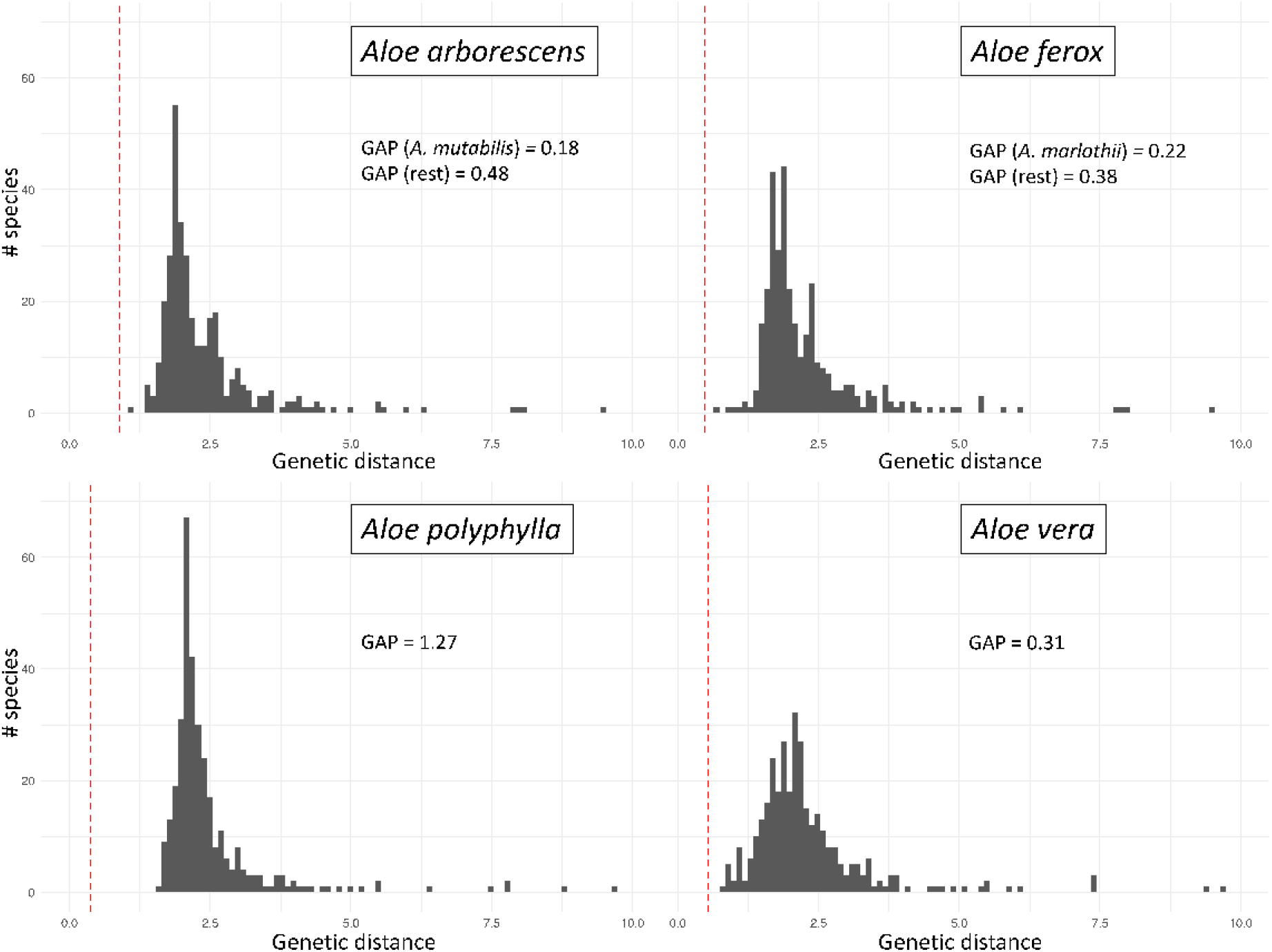
DNA barcoding gaps for four economically important aloes, as determined by the combined pairwise genetic distances for 173 low-copy nuclear loci (section 1.7 of Supporting Information). Distances (defined as the number of nucleotide substitutions per 1000 bp) are relative to the morphologically verified reference dataset sample of each respective species. The DNA barcoding gap is calculated as the difference between the lowest interspecific distance (leftmost dark grey bar) and the highest intraspecific distance (red dotted lines). For *Aloe arborescens* and *A. ferox* two gaps are indicated: one with the most closely related species that is also debated as potentially conspecific and one with all the other species (considering the debated species as conspecific).

### 4. Conservation target assessment

More than a third of all aloes (213/616 species) are threatened with extinction, representing 49% of all assessed species (Figure 1B). This follows from our consensus extinction risk assessment dataset in which we complemented 282 IUCN Red List assessments^23^ with 93 pending IUCN Red List assessments for Madagascar (prepared by one of us: SR) and 157 national red list assessments for South Africa (SANBI Red List 2024: http://redlist.sanbi.org/). Most of these threatened species are endemic to Madagascar (95 species), where 82 species are endangered, of which 23 in critical condition. One of these species, *Aloe fragilis*, was discovered among the intercepted samples (Table 1, “CITES-05-1”; Figure 3E). Another sample belonged to the *Aloe somaliensis* complex from the Horn of Africa, where three out of five species are facing severe population decline (Figure 3). The rest of the intercepted samples all belong to South African species, two of which are “near threatened” (*Aloe arenicola* and *Aloe reynoldsii*). Two samples identified as *Aloe ferox*, for which finished products are now exempt from CITES regulations, and two as *Aloe polyphylla*, for which no extinction risk assessments are available.

## Discussion

The Anthropocene presents dire conditions for wild species, with 28% of all assessed species threatened with extinction^23^. With 49%, the aloes are far above this and can thus be considered one of the most endangered plant groups on the planet. Coupled with high economic interest and a realistic threat of wild overharvesting, an accurate identification tool is both timely and critical to monitor international trade. Using a state-of-the-art molecular identification technique, we present a reliable solution to DNA barcoding of *Aloe vera* and endangered relatives. Our protocol generates high-coverage DNA sequence data for 189 independently evolving nuclear genes, presenting well established DNA barcoding gaps for economically important species - and therefore confident identification - while providing a clear indication of uncertainty due to taxonomic complexity (if present) through gene tree conflict. The largest-to-date phylogenomic effort for *Aloe* and related genera, comprising >300 species enables accurate identification of aloe leaf samples that would otherwise be impossible to identify even by expert taxonomists. With this tool we successfully identified confiscated samples to species level, presenting a significant innovation beyond previous identification protocols. Previous efforts relied on diagnostic morphological characters^19^ - that are rarely preserved in semi-processed trade samples - and uninformative, highly conserved organellar DNA markers^8,21^. Barcode-based diagnostics are well established for timber products (e.g., mahogany and laurel)^8^ and cycads^4^, whereas aloes have been notoriously difficult to barcode, and success has been limited using classic barcode markers^9^, random amplified polymorphic DNA^28^, and full plastid genomes^21^.

The use of customised target capture sequencing^15^ takes advantage of more variable nuclear genomic loci and works efficiently on degraded DNA typical of processed and museum specimens^16^. This is due to the small size of oligonucleotide probes^14^ and enabled us to include historic DNA from 51 herbarium specimens. The new barcoding tool is applicable to a wide range of species, as compared to authentication techniques such as chemical fingerprinting which achieve high resolution but tend to be narrowly focused on few species (e.g. *Aloe ferox*^29^). While the benefits of DNA-based diagnostics are well established, successful protocols are contingent on comprehensive and taxonomically verified reference sequence datasets. Each reference sequence must be curated with an accompanying herbarium voucher specimen^30^ providing the physical record of the source biological material from which the DNA sequence is derived. With the use of botanical collections, reference material comes directly from a verified voucher specimen, and this has, as such, enabled the (relatively rapid) generation of a reference sequence database with broad taxonomic coverage in aloes.

Ease of application positions DNA barcoding as a near real-time monitoring tool for conservation and trade, and may reveal contemporary targets for international conservation action. Illegal plant trafficking (IPT), such as succulent poaching, is an increasingly worrying threat to many plant species^27,31,32^. In wildlife source areas, such as sub-Saharan Africa, IPT impacts are exacerbated by increased online demand^2^ coupled with continued natural habitat encroachment^31^.

The necessity for more diagnostic power in diverse and recently evolved clades, such as aloes^25^, has now brought DNA barcoding into the nuclear genomic era^13,14^. When verified and accurate, these tools can offer tremendous progress in the monitoring of wildlife trade and help reveal unsustainable practices^33,34^. Similarly, these tools can be deployed by producers of biological products to substantiate their sustainable produce^35^. The informative low-copy nuclear genes captured with this tool^15^ were previously inaccessible due to the high cost of generating high-coverage sequence data on such large genomes as found in aloes (15 Gb on average^36^). With target capture, this problem is circumvented as it reduces the complexity of genomic sequence libraries to target loci, regardless of genome size. Applying this tool to a variety of trade samples, we found evidence of legitimacy in the aloe leaf market, but also revealed concerns in the movement of CITES-regulated species. Considering the improvements in aloe identification in our study, we encourage the use of target capture sequencing for other diverse groups subject to IPT, such as orchids^37^ and cacti.

Extinction risk for aloes is higher than the background rate of half of all plant species^38^ (Figure 1 & 3). In Madagascar, an important centre of diversity with 142 species of aloes^22^, two thirds of the assessed species are threatened with extinction (Figure 1B), comparable to cycads^23^ including 23 species that are endangered (online supporting material). Although land use change (e.g. agriculture, mining) is the main threat for these species^19^, IPT for horticulture potentially presents a significant additional pressure. The identification of *Aloe fragilis*, a horticulturally valued but critically endangered species listed on CITES appendix I, among the Heathrow interceptions is therefore worrying.

Plant biodiversity in Southern Africa is strongly correlated with human population density, putting a tension between people and plant conservation^39^. Aloes can be local targets for bioremedies and food^40^ and the use of (potentially toxic) substitute species compromise health as well as local biodiversity. The DNA barcoding tool could be used to correctly identify species in use, monitor trade in the two cultivated non-CITES species *Aloe vera* and *A. ferox*, and identify CITES-regulated aloe species in burgeoning substitute industries^18^ for them.

Finally, we note that the conservation impact of this study relies, in a large part, on accurate and up-to-data extinction risk assessments. There are considerable gaps in this dataset for aloes, particularly for species from the Arabian Peninsula, where only three (out of 50) species have been properly assessed (Figure 1). For Madagascar, we could update the dataset considerably with new assessments by local botanists, bringing the coverage up to 85%. The gap in official IUCN Red List assessments is also severe for South Africa (138 species not assessed or lacking data), but we were able to complement this with a near-complete national red list (SANBI)^24^, highlighting the importance of local conservation initiatives. Conservation situation can change very rapidly for species, highlighted by considerable differences we found in our new assessments (for Madagascar) and national red list data (for South Africa) compared to the official IUCN Red List data (online supporting material). The Madagascan *Aloe delicatifolia*, for example, is officially listed as Least Concern^23^ but updated to Endangered in our new assessment. As we show here, extinction risk data and DNA barcoding tools can work in harmony to highlight specific international conservation targets related to IPT. We hope that our example will inspire international and local colleagues to implement action of both sides of this interaction to help safeguard threatened plant species in the Anthropocene.

## Acknowledgements

The authors express gratitude to all the horticultural and herbarium curatorial staff for their efforts in generating, maintaining, and curating the botanical collections at the following institutes: Royal Botanic Gardens, Kew, and Kew Herbarium (K); National Museums, Tropical East Africa Herbarium (EA); Uppsala Botanic Garden and University of Uppsala Herbarium (UPS); Gothenburg Botanic Garden; Potsdam Botanic Garden; Musée National d’Histoire Naturelle, Paris (P); Cambridge University Botanic Garden. We thank Anne-Sophie Quatela for advice on DNA extractions from herbarium material. Computational resources were provided by UNINETT Sigma2 - the National Infrastructure for High Performance Computing and Data Storage in Norway.

## [DATA AVAILABILITY STATEMENT]

Raw sequencing data newly generated for this study to produce the Alooideae reference dataset and identify unknown aloe DNA samples are deposited in the Sequence Read Archive (SRA) of the U.S. National Center for Biotechnology Information (NCBI) under Bioproject PRJNA1120847 (for “aloes”) and under Bioproject PRJNA1122593 (for non-aloes). Supplementary sequencing data from the Alooideae target capture tool design study (Woudstra et al. 2021, Scientific Reports) is available in the SRA at NCBI under Bioproject PRJNA1120785. Other data related to this manuscript, including the reference sequences for the four species used in the design of the Alooideae target capture tool are available through online supporting material deposited in FigShare at DOI: 10.6084/m9.figshare.24487750.

## [SUPPORTING INFORMATION]

A detailed description of the methodologies and results presented in this study is available in the Supporting Information.

## [FUNDING STATEMENT]

This study was funded by the European Union’s Horizon 2020 framework, as part of the MSCA-ITN-ETN Plant.ID under agreement number 765000.

## [CONFLICT OF INTEREST DISCLOSURE]

The authors declare to have no conflicts of interest related to the study.

## [ETHICS APPROVAL STATEMENT]

Not applicable.

## [PERMISSION TO REPRODUCE MATERIAL FROM OTHER SOURCES]

Not applicable.

## Notes

https://doi.org/10.6084/m9.figshare.24487750

### Competing Interest Statement

The authors have declared no competing interest.

