## Supporting Information for "An updated DNA barcoding tool for *Aloe vera* and related CITES-regulated species"

*Supplemental to Woudstra, Y, et al. 2024 - An updated DNA barcoding tool for Aloe vera and CITES-restricted relatives*

Data and scripts related to this manuscript are deposited in the FigShare repository at DOI: 10.6084/m9.figshare.24487750

1. Alooideae target capture tool reference sequences ("Aloeref.fasta")
2. Accession information file
3. Sequencing information file
4. Extinction risk & geographical distribution information file
5. R script for producing Figure 1 (of main manuscript)
6. Data file to produce plots for Figure 1 ("IUCN_renumbered.csv")
7. Co-phylogeny of reference database topologies from concatenation vs. coalescent-based methodologies
8. Phylogeny with distribution of CITES-appendix listings

Raw sequencing data are deposited in the NCBI short read archive (SRA) consists of the following bioprojects:
PRJNA1120785: Alooideae target capture design – pilot study
PRJNA1120847: Alooideae reference database (aloes) & identifications of unknown aloe material
PRJNA1122593: Alooideae reference database (non-aloes)

### Part 1: Detailed methods overview

#### Materials & methods

*1.1 Sampling and plant material*

The group of plants collectively called "aloes" comprises the diverse genus *Aloe* (595 species and 55 additional subtaxa), the smaller genera *Aloiampelos* (7 species and 4 additional subtaxa), *Aloidendron* (6 species), *Gonialoe* (4 species) and *Kumara* (2 species), and the monospecific genera *Aloestrela* and *Aristaloe*. For building a comprehensive reference database, a sampling strategy was designed to accommodate all genera as well as taxonomic groups described in monographs[1–4], phylogenetic clades[5–7] and geographic centres of diversity[8] for the complex genus *Aloe*. Samples were obtained either from living collections in botanic gardens or from herbarium collections and supplemented by samples used in previous studies. Detailed information on sample origins can be found in the accession information file (online supporting material).

375 samples were prepared belonging to the genus *Aloe*, representing 352 species, four additional subspecies and eight additional varieties. An additional ten species belonging to other genera comprising the "aloes" were sampled, including *Aloestrela*[9] (one species), *Aloidendron* (four species)[10], *Aloiampelos* (two species)[10], *Aristaloe* (one species), *Gonialoe* (one species)[5] and *Kumara* Medik. (one species)[10]. To obtain a good representation of the remaining members of the Alooideae subfamily, we included one species each of the genera *Astroloba*[11], *Haworthia*[12], *Haworthiopsis* and *Tulista*[5].

Freshly harvested leaf material was processed into ±1 $cm^2$ pieces of leaf tissue, with the clear layer of the leaf mesophyll removed, and dried in silica powder with indicator beads for at least one week. For dried and pressed herbarium vouchers, similarly sized leaf fragments or dried flowers were taken from the specimen. Destructive sampling of type specimens was carried out for some species (sequencing information file, online supporting material) and only when this was not destroying important characters for identifying the species, with permission from the relevant curator.

The abovementioned samples were combined with published sequences for 29 species, used in the Alooideae bait panel design pilot study[13]. These comprised sequences for 24 Alooideae species obtained through target capture (23 *Aloe* species and one *Aloiampelos*) and for four Alooideae species obtained with transcriptomes generated to design the Alooideae bait panel (three *Aloe* species and one *Aloidendron*). We used sequences from *Bulbine frutescens* (subfamily Asphodeloideae), obtained with the same target capture tool, as an outgroup to the Alooideae clade.

*1.2 DNA isolation and purification*

Dry samples of leaves or flowers (c. 20 mg) were ground into a fine powder using stainless steel beads in 2mL Eppendorf tubes using a TissueLyser. For silica dried leaf samples, DNA was isolated using a Qiagen DNEasy® extraction kit (Qiagen, Hilden, Germany) following the manufacturer's instructions. Tissue from herbarium specimens, in which DNA can be degraded by the age of the specimen, drying method or other treatments[14–16], was treated with a CTAB protocol[17,18] to obtain higher yields. Some samples were first treated with a wash step involving a sucrose-tris-EDTA (STE) buffer to remove excess polysaccharides[19] (sequencing information file).

Total genomic DNA isolated with the CTAB method was subsequently treated with AMPure XP beads (Beckman Coulter, Brea, California, USA) for purification. Bead-bound DNA was precipitated on a magnetic tube rack and washed 2-3 times with freshly prepared 70% ethanol, after which DNA was eluted from the beads in 10 mM tris-EDTA (TE). In some cases, DNA was eluted several times from the beads to improve yield and subsequently concentrated by vacuum evaporation. DNA isolated with the Qiagen DNEasy® kit was not further purified since the isolation protocol contains a purification step using silica-columns, and no significant impurities were evident in the fluorescence spectrum of the DNA isolated with this method (e.g., $Abs_{260/230}$ ±2.4 and $Abs_{260/280}$ ±1.8).

The concentration of DNA was quantified with a Quantus™ fluorometer (Promega, Maddison, Wisconsin, USA) and DNA fragment size was estimated either by agarose gel electrophoresis or by Genomic DNA ScreenTape analysis on an Agilent 4200 TapeStation system (Agilent, Santa Clara, California, USA).

*1.3 HTS library preparation*

High-molecular-weight samples (accession & sequencing information file) were fragmented to the appropriate insert size by ultrasonication in 50 µL TE buffer in 130 µL Covaris® tubes using an M220 Focused Ultrasonicator (Covaris, Woburn, Massachusetts, USA). HTS libraries were created using the NEBNext® Ultra™ II DNA library preparation kit with accompanying Multiplex Oligos Dual Index sets 1&2 (New England Biolabs, Ipswich, Massachusetts, USA) for short-read paired-end sequencing on Illumina® HiSeq X (2x150bp) platforms (Illumina, San Diego, California, USA) with a starting quantity of ≤100ng DNA (sequencing information file). Library quality was assessed on DNA quantity and fragment length analysis on the Quantus™ fluorometer and by D1000 ScreenTape on an Agilent 4200 TapeStation, respectively.

*1.4 Pooling and nuclear target enrichment*

Libraries were diluted to 10nM concentration and pooled in batches of 8-27 equimolar samples, based on either taxonomic affinity, library fragment size or a combination of both (sequencing information file), following recommendations in the literature [20,21]. Using a custom-designed target capture RNA-bait panel for Alooideae[13], each pool was hybridised with one reaction of the specific myBaits® kit (Arbor Biosciences, Ann Arbor, Michigan, USA) following the manufacturer's protocol (myBaits® manual v3). Hybridisation of bait-target DNA complex was performed at 65°C for approximately 24 hours and washing was performed in 1.5 mL DNA LoBind Eppendorf tubes. Some pools were enriched with half-volume reactions of baits (sequencing information file). Post-capture amplification was performed with universal P5 and P7 primers, complimentary to the NEBNext® adaptors for Illumina® sequencing, using 18 cycles with optimised extension time for the expected enriched library fragment size, following the myBaits® protocol.

*1.5 Sequencing, read trimming and target assembly*

Enriched pools were diluted to 10 nM concentration, based on pool DNA concentration and fragment size distribution measurements (as above), and combined in equimolar concentrations for sequencing on the Illumina HiSeq X platform. Sequencing was outsourced to Macrogen Inc. (Seoul, Republic of Korea). 150 bp paired-end reads were generated by sequencing up to 190 samples per lane with compatible dual index barcodes. Raw-read data was retrieved after demultiplexing by Macrogen Inc. using Illumina's BaseSpace software.

The quality of the raw reads was assessed using FastQC[22], after which the reads were trimmed using Trimmomatic v0.39 (Bolger, Lohse, and Usadel 2014) to remove terminal bases with a phred-33 score below 30 (e.g., <99.9% base call accuracy). Trimmed reads were then assembled into target loci using the HybPiper v1.3.1[23] pipeline with the sequences used in the *Aloe* bait panel design[13] as the assembly reference. Assembled target loci were retrieved as exon-only (pipeline default output) for optimal sequence recovery overlap between samples. HybPiper can also extract sequences belonging to flanking intronic and intergenic regions from (overhanging) off-target reads, which are generally more variable between species than exonic regions[24]. However, we chose not to pursue this option for our molecular identification study because of the relative randomness in recovery for these regions due to the *Aloe* baits[13] being designed for exonic regions only. All loci were checked for potential paralogy using the paralog warning script in HybPiper. For downstream phylogenetic analysis, only samples with a target recovery of ≥50% of the total target length (≥175 kbp) were retained.

*1.6 Target sequence alignment and phylogenetic tree estimation*

To produce a reference database for the molecular identification exercise, assembled target exon sequences were aligned per locus into MSAs using MAFFT v7.429[25], which were cleaned in three steps after removing paralogous (section 1.5) and poorly recovered loci (<200 samples). First, CIAlign v1.0.14[26] was called to remove all insertions (<650 bp) and deletions present in <50% of the samples and to remove sequences <100 bp long. Second, Phyutility v2.2[27] was called to remove all columns covered by <20% of the samples. Finally, manual cleaning was performed by visualisation in Geneious 9.1 Pro (http://www.geneious.com/) to remove previously undiscovered putative paralogous (regions of) loci and poorly recovered insertions not removed by CIAlign and Phyutility.

Two methods were employed for phylogenomic tree estimation: a maximum likelihood approach using the concatenated supermatrix of all MSAs remaining after cleaning, and a coalescent-based approach in which incongruence between gene trees can be estimated as

well. A supermatrix was built by concatenating all MSAs after cleaning, using FasConCAT-G v1.04[28]. Maximum likelihood trees were estimated both from the supermatrix to produce a species tree and from the individual-locus MSAs to produce gene trees using IQTree v1.6.12[29]. We used a general time reversible (GTR) model combined with a gamma-distribution for rate heterogeneity and a correction for invariable sites (GTR+G+I settings). Support for the nodes in the consensus trees was calculated using 1000 ultra-fast bootstrap replications.
A coalescent-based species tree was estimated from the gene trees with ASTRAL-III[30]. Incomplete lineage sorting was estimated by gene tree discordance, reported by ASTRAL-III as the ratio of quartets that are shared between all gene trees and the species tree. To determine incongruence between the concatenated and coalescent-based approaches, we compared the two topologies in a cophylogeny in R v4.0.3[31] using the cophylo function from the package phytools v0.7-70[32]. Trees were visualised in the online Interactive Tree of Life (iTOL) tool[33], rooted on the branch containing our outgroup sample of *Bulbine frutescens*, and annotated using the iTOL annotation editor.

*1.7 Customised DNA barcoding*
To test the sensitivity of the custom *Aloe* bait panel as a DNA barcoding tool, we attempted to enrich and sequence 189 low-copy nuclear genes in 39 samples with unknown or uncertain identity. Identification was done through phylogenomic inference, comparing a concatenation-based method to a coalescent-based method (section 1.6), contrasted with a more traditional method using pairwise genetic distances.
To validate the protocol, one of us (PR) prepared ten anonymised samples from previously unsampled accessions from the living collections at Royal Botanic Gardens, Kew (RBG Kew). These accessions had verified taxonomic identities based on flowering morphology and provenance, providing a solid identity to test the tool on. The validated tool was then applied in a conservation context with ten samples of CITES-restricted plants (Figure 2 of main manuscript) seized by border control at London Heathrow Airport that were sent to RBG Kew, the UK CITES Flora authority, for identification. Additionally, we obtained three fresh leaf samples from the UK domestic and international market to test the tool on traded samples. Two samples were sold as *Aloe vera* leaves from organic supermarkets in London, United Kingdom and the third one was sold as *Aloe arborescens* Mill. in a homoeopathic medicinal package sold by an Italian farm. Finally, we applied the tool to the curation of unidentified specimens at RBG Kew using six unlabelled plants in the living collections and ten unidentified wild-collected herbarium specimens.
DNA was extracted from silica dried and herbarium voucher specimens using the protocols described above (section 1.2). One herbarium voucher did not yield enough clean DNA (<1 ng) to be considered for HTS library preparation. The rest of the DNA samples were fragmented to the appropriate size (if necessary), prepared for HTS sequencing and enriched with the *Aloe* bait panel as described above (sections 1.3 and 1.4). Sequencing, read trimming and target assembly was performed in parallel with the taxonomic samples intended for the reference database (section 1.5). Like with building the reference database, we filtered out samples with a total target recovery below 50% from the molecular identification exercise.
Identifications were performed with three different methods: the concatenation- and coalescent-based phylogenomic approaches (section 1.6) and a pairwise genetic distance approach. The identification algorithm was performed twice. First for the ten verified and anonymised accessions to validate the tool, after which the reference database was updated. Second for all the unidentified samples combined. We combined the low-copy nuclear target

sequences of the unknown samples with the reference database, re-aligned MSAs for each locus and followed the MSA cleaning protocol in section 1.6. Phylogenomic inferences were then repeated for the updated dataset and sample identities were determined as the taxon with which the unknown sample is recovered in a monophyletic clade, with the node support value indicating the confidence in this identity. In the case of non-monophyly, the closest relative of the unknown sample was noted, if applicable.

A pairwise genetic distance matrix was generated using the function distmat of the command-line package EMBOSS v6.6.0[34]. We used the uncorrected substitution algorithm in which the programme simply counts the number of substitutions and penalises for gaps, with the final score representing the number of substitutions per 1000 bp. Sample identities were determined as the taxon with the lowest pairwise genetic distance, recording the distance as an indication for the closeness to the reference taxon. Additionally, we recorded the gap between the intra- and interspecific genetic distances (the so-called 'barcoding gap'[35]) as a proxy for identification confidence.

*1.8 Conservation target assessment*

Identifying the *Aloe* species among CITES interceptions could help identify conservation targets. To make the correlation between our identifications and conservation status, we updated extinction risk assessments among the aloes and plotted these on the phylogenetic framework obtained with our reference database. We downloaded the current publicly available IUCN Red List assessments (release 2023-1) from the official website (https://www.iucnredlist.org) and partially filled gaps in this dataset with new unpublished (pending) assessments from Madagascar (provided by SR) and published data from the South African National Biodiversity Institute (SANBI) Red List of South African Plants (http://redlist.sanbi.org/index.php). SANBI uses the same criteria as the IUCN Red List, except for an extra category called "Rare" meaning a species is highly localised with very few sightings. SANBI treats this category in a comparable way to the IUCN Red List's "Near Threatened" (NT) category (http://redlist.sanbi.org/stats.php) and SANBI data for aloes in our dataset was thus converted to this accordingly. For three species (*A. nubigena*, *A. pearsonii* and *A. peglerae*) the SANBI and IUCN datasets disagreed on the extinction risk, and we retained the highest risk assessment (extinction risk and geographic distribution information file). The distribution of extinction risk for the main geographic regions of Alooideae was analysed in a bar plot in R using ggplot2 v3.3.5[36] (R script for producing Figure 1). Regions were defined to represent main centres of diversity for the genus *Aloe*[6]: Arabian Peninsula (AP); Horn of Africa (HA); Madagascar (MA); Namibia (NM); Southern Africa (SA); Tropical East Africa (EA); West Africa (WA); South Tropical Africa (ZA).

These assessments were plotted on the coalescent-based phylogeny for the updated Alooideae reference database obtained after the validation of the molecular identification tool (section 1.7) using iTOL annotation editor[33]. Species identified among our CITES-restricted samples were then manually highlighted on the tree to visualise the correlation with extinction risk.

Although all members of the *Aloe* genus (including those previously belonging to *Aloe*), except *Aloe vera* and *Aloe ferox* are protected under CITES legislation, transportation for some species is more heavily restricted than for others. If a taxon is included under Appendix I all international transportation is restricted, if included in Appendix II international transportation is allowed but permits are required. To see the phylogenetic distribution of these different levels of CITES protection, we repeated the above process used for extinction risk assessment with CITES appendix listing data (https://cites.org/eng/app/appendices.php).

**Part 2: Detailed results overview**

*2.1 Target capture sequencing*

The number of paired reads (150 bp each) after trimming and filtering ranged from 1,981 to 61,398,381 (average 4,304,079) with an average of 23.8% of the reads on-target. Target recovery was successful (>50% of the total target length) for 352 samples (>74% of 474 samples in total). Among the approved samples, the average target recovery was 304,858 bp (87.5% of the total target length) with the number of target loci with sequences ranging from 172 to 189 (the total number of loci targeted). Considering all samples, differences in target recovery were negligible between lower (≤ 24 samples to 1 reaction) and higher (≥42 samples to 1 reaction) pooling strategies, with 66.0% compared to 66.3%, respectively.

DNA samples fitted into three categories according to the origin of the material: DNA stocks from previous studies (>5 years old, varying degrees of degradation, 70 samples), DNA extracts from newly collected silica-dried material (average fragment length >10 kbp, 244 samples) and DNA extracts from herbarium specimens (average fragments generally <1 kbp, 156 samples). Target recovery was considerably higher from non-ancient DNA, with an average of 89.5% for DNA stocks and 81.2% for silica-dried material, compared to 31.5% in herbarium material. Herbarium samples therefore accounted for the majority (105/122) of failed samples in the study. Ancient DNA (aDNA) isolated with the STE-pre-treatment did not perform better than aDNA obtained from the regular CTAB method, with an average of 22.9% target recovery and a 50% failure rate, compared to 32.2% target recovery and a 53% failure rate for those not treated with STE. 42 samples had a target recovery of >10,000 bp, despite having >100,000 reads on-target. This was caused by several baits aligning with highly repetitive elements in some samples, as revealed by visual inspection of the HybPiper mapping results.

Detailed target capture sequencing results per sample are listed in the accession & sequencing information file.

*2.2 Alooideae reference database*

The reference dataset for Alooideae consisted of DNA sequences obtained from 310 samples representing 285 species of *Aloe* (including 5 subspecies and 4 varieties), 4 species of *Aloidendron*, 3 species of *Aloiampelos* and 1 species each of the genera *Aloestrela, Aristaloe*, *Astroloba*, *Haworthia*, *Haworthiopsis*, *Gasteria*, *Gonialoe*, *Kumara* and *Tulista*. Additionally, outgroup taxon *Bulbine frutescens* was included for proper rooting of the trees. The full dataset was trimmed from 189 target loci to 173, after removing paralog-containing alignments and poorly recovered loci (<200 samples represented). The obtained topologies recovered the same relationships between Alooideae genera.

There were, however, considerable differences between the coalescent and maximum likelihood topologies in the placement of taxa sharing the MRCA with all *Aloe* species (e.g., clades involving *Aloe polyphylla*, *A. comptonii*, *A. microstigma* clade, and *A. arborescens*), indicating incomplete lineage sorting for the genus *Aloe*. This was also reflected in the normalised quartet score of 0.622 obtained with ASTRAL-III for the coalescent topology. Most of the topology, however, was consistent in both estimation methods, with large monophyletic clades showing strong geographical patterns (e.g., Namibia, Zambezia (South Tropical Africa), Tropical East Africa, Horn of Africa / Arabian Peninsula, and Madagascar).

Both topologies are available as a co-phylogeny in the online supplemental, indicating topological differences between the coalescent and concatenation methods.

*2.3 DNA barcoding*

Prior to a conservation-minded DNA barcoding exercise, we validated our customised tool with ten anonymised curated samples with morphologically verified identifications. Subsequent sampling for the DNA barcoding exercise consisted of 26 accessions, comprising three samples of *Aloe* leaves bought on the domestic UK and international market, ten samples intercepted by border control at London Heathrow Airport under CITES legislation, six unidentified nursery samples from the living collections at Royal Botanic Gardens, Kew, and nine unidentified herbarium specimens from the Herbarium Kewensis. Only one herbarium sample passed our target recovery filtering (≥ 50% total target) and the others were therefore not considered in the DNA barcoding exercise.

*2.3.1 Validation with anonymised verified samples*

The sensitivity of our custom bait panel and phylogenomic assessment in identifying *Aloe* DNA extracts to species rank was confirmed by the accurate species placement of 60% of anonymised samples ("anonymous-" in Table 1) using the coalescent phylogenomic method (Figure 3). Furthermore, all ten samples were identified to the correct taxonomic clade, thereby allowing confident separation between CITES non-regulated (*Aloe vera* and *A. ferox*) and regulated species (all others). Three of the ambiguous samples (Anonymous-03, -04 and -06) belonged to species that were not yet present in our reference database (*A. pendens*, *A. millotii* and *A. lineata*, respectively) and were henceforth added to update the database. The only true misidentification, therefore, was in the *Aloe somaliensis* species complex, where "Anonymous-07" was morphologically verified as *A. jucunda* but identified as *Aloe somaliensis* with DNA barcoding. The genetic distance analysis indicated *A. jucunda* as the likely origin of this material, but the low barcoding gap (0.05 substitutions per 1000 bp) gave evidence for poor separation between species in this complex. "Anonymous-02", on the other hand, belongs to the same species complex but was confidently assigned to *A. peckii*. Gene tree conflict is high in this clade, as indicated by the moderate support (LPP=0.71) for the misidentification of "Anonymous-07" and for the separation between *A. jucunda* and *A. somaliensis* (LPP=0.61).

A lower success rate of 50% species rank identification was observed using the concatenation phylogenomic and genetic distance methods. All three versions of our tool gave a different placement to sample Anonymous-03, with the support value obtained with the moderate support in the coalescent phylogenomic method (LPP=0.53) indicating high levels of gene tree incongruence in the species complex of Yemeni aloes. "Anonymous-09" belongs to the species complex involving *A. pachygaster* and *A. claviflora*. The coalescent phylogenomic method yielded an equal probability for these two species, whereas pairwise genetic distances favoured *A. pachygaster* as the source species. The concatenated phylogenomic method incorrectly indicated the less related *A. hereroensis*.

We therefore considered the coalescent phylogenomic version of our DNA barcoding tool validated on the basis that every sample was assigned to the correct taxonomic, with gene tree conflict and phylogenetic support giving clear and reliable indications of caution.

*2.3.2 International market samples*

All versions of the tool identified the three market samples (Figure 4) to the species indicated by the seller, although it was not possible to discount *A. mutabilis* as a potential alternative to *A. arborescens* for sample Market-1 (Table 1). This was indicated by an undecided identification using the concatenated phylogenomic version, moderate support (0.77) using the coalescent phylogenomic version, and a low barcoding gap (0.13, Figure 5A) using the genetic distance version.

*2.3.3 Plant material seized by law enforcement*

Among intercepted illegal trade material, identifications were highly supported and fully congruent between the two phylogenomic versions of the tool. These interceptions were non-randomly distributed in terms of phylogenomic and likely geographic origins (Figure 4), with 8/10 samples belonging to South African species. In *Aloe ferox* and *A. polyphylla*, we discovered two species that were represented by two illegally traded samples. Apart from *A. polyphylla*, for which the extinction risk is unknown, none of the South African samples seem to belong to threatened species. However, the one Malagasy sample among the interceptions belongs to the critically endangered *A. fragilis*. The final sample likely originated from the Horn of Africa, given its affinity with the *A. somaliensis* species clade, in which 4/6 species are threatened.

Two samples (CITES-08 and -09) had enigmatic identities, both of which seem to belong to species not sampled for our reference database. The two identifications were not fully supported using the coalescent phylogenomic version (LPP=0.90 and 0.91, respectively), indicating some gene tree incongruence. Furthermore, pairwise genetic distances with the closest reference sample were high (>1.2), while the barcoding gap was very low (<0.05). Sample “CITES-08” is a member of the *A. somaliensis* species complex and “CITES-09” is part of a clade comprising *A. vryheidensis*, *A. alooides* and *A. globuligemma*. In the diverse clade of spotted (maculate) aloes (*Aloe* sect. *Pictae*) results with genetic distance diverged from phylogenomic identifications for two samples (CITES-02 and -10) where the barcoding gap was low (0.06 and 0.02, respectively).

*2.3.4 Unlabelled specimens from botanical collections*

The customised *Aloe* DNA barcoding tool showed encouraging results when applied to help verify specimens for curation in the Living Collections and Herbarium at the Royal Botanic Gardens, Kew. Four living plant specimens were identified to species rank with high confidence according to their placement in both the coalescent and maximum likelihood trees (Table 1, Figure 3). The two remaining living plant specimens received conflicting identifications between the different versions of the DNA barcoding tool, with very low support using the coalescent phylogenomic method and very small barcoding gaps (<0.10).

Identification of the herbarium specimens was less successful. Only one indeterminate herbarium sample passed the target recovery filtering criteria (≥50%), belonging to a specimen originally collected on St. Helena Island as *Aloe vera*. Both phylogenomic methods of our tool fully discarded this identity with strong support (LPP=0.99, BS=100). Using the concatenation result, this sample could be placed in the maculate aloes, whereas the coalescent version of our tool even placed this sample on a lone branch. The genetic distance with the nearest *Aloe* reference sample was also very high at 2.57.

*2.3.5 Establishing DNA barcoding gaps*

DNA barcoding gaps were analysed in detail for four *Aloe* species with considerable economic interest (Figure 5), to which we identified one or more samples using our DNA barcoding exercise. For the South African *Aloe arborescens* and *A. ferox*, poorly distinguishable species in previous molecular phylogenetic studies, barcoding gaps were small (<0.25) due to potentially conspecific taxa present in the reference database (*A. mutabilis* and *A. marlothii*, respectively). Considering these taxa conspecific considerably increased the barcoding gap with 167% and 72%, respectively. For *Aloe vera*, a decent barcoding gap of 0.31 existed, despite this taxon being nested in a phylogenomically poorly resolved clade. The final barcoding gap (considering potentially conspecific taxa) decreased with phylogenetic distance from the MRCA of all aloes, from 1.27 for *A. polyphylla* to 0.48 for *A. arborescens*, to 0.38 for *A. ferox*, and finally to 0.31 for *A. vera*. This also reflects the number of closely related sister taxa in a clade which increases from 0 for *A. polyphylla*, to 6 for *A. arborescens*, to 7 for *A. ferox* and finally to 18 for *A. vera*.

*2.4 Extinction risk distribution in the Alooideae*

IUCN Red List assessments were available for 298 aloes, comprising 290 *Aloe* species, one *Aloestrela*, one *Aloiampelos*, four *Aloidendron* and one *Gonialoe*. Fourteen of these assessments were data deficient (DD), the majority of which are from the Horn of Africa. The largest knowledge gap is in the Arabian Peninsula, where only three out of fifty species have been properly assessed. A list of 93 pending assessments (provided by SR) considerably filled gaps for Madagascar, pushing the total up to 123 species assessments for this region. A national red list for South Africa (SANBI Red List of South African Plants), comprising 146 *Aloe*, nine *Aloiampelos* and five *Aloidendron* assessments, filled gaps to a near complete coverage (95%) of extinction risk assessments for that region.

Extinction risk is high for the aloes, with one in three aloes (212 species) being threatened (categories VU, EN, CR; Figure 1B). The rate of threatened species is lowest for South African (37/148 assessments) and Namibian (1/18) species, but very high for Madagascar (94/122) where one species (*Aloe silicicola*) is considered extinct in the wild. This is reflected in the phylogenetic distribution of extinction risk (Figure 3), as well as CITES appendix 1 listings where the majority (18/21, Phylogeny with distribution of CITES-appendix listings; online supporting material) is of Malagasy origin. The rate of threatened taxa was also high in the Tropical East Africa region (31/51), but lower in the Horn of Africa (34/87) and South Tropical Africa (13/37) regions. The only described species from India (*Aloe trinervis*), on the other hand, is an endangered plant.

2010).

16. Forrest, L. L. *et al.* The Limits of Hyb-Seq for Herbarium Specimens: Impact of Preservation Techniques. *Front. Ecol. Evol.* **7**, 439 (2019).
17. Doyle, J. J. & Doyle, J. L. A rapid DNA isolation procedure for small quantities of fresh leaf tissue. *Phytochem. Bull.* **19**, 11–15 (1987).
18. Doyle, J. J. & Dickson, E. E. Preservation of Plant Samples for Dna Restriction Endonuclease Analysis. *TAXON* **36**, 715–722 (1987).
19. Shepherd, L. D. & McLay, T. G. B. Two micro-scale protocols for the isolation of DNA from polysaccharide-rich plant tissue. *J. Plant Res.* **124**, 311–314 (2011).
20. Hale, H., Gardner, E. M., Viruel, J., Pokorny, L. & Johnson, M. G. Strategies for reducing per-sample costs in target capture sequencing for phylogenomics and population genomics in plants. *Appl. Plant Sci.* **8**, (2020).
21. Woudstra, Y. *et al.* Target Capture.. in *Molecular Identification of Plants: From Sequence to Species* (Pensoft Publishers, 2022).
22. Andrews, S. FastQC: A Quality Control tool for High Throughput Sequence Data. (2010).
23. Johnson, M. G. *et al.* HybPiper: Extracting coding sequence and introns for phylogenetics from high-throughput sequencing reads using target enrichment. *Appl. Plant Sci.* **4**, 1600016 (2016).
24. Villaverde, T. *et al.* Bridging the micro- and macroevolutionary levels in phylogenomics: Hyb-Seq solves relationships from populations to species and above. *New Phytol.* **220**, 636–650 (2018).
25. Katoh, K., Standley, D.M. MAFFT Multiple Sequence Alignment Software Version 7: Improvements in Performance and Usability. *Mol Biol Evol.* **30**, 772-780 (2013).
26. Tumescheit, C., Firth, A. E. & Brown, K. CIAlign - A Highly Customisable Command Line Tool to Clean, Interpret and Visualise Multiple Sequence Alignments. http://biorxiv.org/lookup/doi/10.1101/2020.09.14.291484 (2020) doi:10.1101/2020.09.14.291484.
27. Smith, S. A. & Dunn, C. W. Phyutility: a phyloinformatics tool for trees, alignments and molecular data. *Bioinformatics* **24**, 715–716 (2008).

28. Kück, P. & Meusemann, K. FASconCAT: Convenient handling of data matrices. *Mol. Phylogenet. Evol.* **56**, 1115–1118 (2010).
29. Nguyen, L.-T., Schmidt, H. A., von Haeseler, A. & Minh, B. Q. IQ-TREE: A Fast and Effective Stochastic Algorithm for Estimating Maximum-Likelihood Phylogenies. *Mol. Biol. Evol.* **32**, 268–274 (2015).
30. Zhang, C., Rabiee, M., Sayyari, E. & Mirarab, S. ASTRAL-III: polynomial time species tree reconstruction from partially resolved gene trees. *BMC Bioinformatics* **19**, 153 (2018).
31. R Core Team. R: A language and environment for statistical computing. R Foundation for Statistical Computing (2021).
32. Revell, L. J. phytools: an R package for phylogenetic comparative biology (and other things): *phytools: R package*. *Methods Ecol. Evol.* **3**, 217–223 (2012).
33. Letunic, I. & Bork, P. Interactive Tree Of Life (iTOL) v5: an online tool for phylogenetic tree display and annotation. *Nucleic Acids Res.* **49**, W293–W296 (2021).
34. Rice, P., Longden, I. & Bleasby, A. EMBOSS: The European Molecular Biology Open Software Suite. *Trends Genet.* **16**, 276–277 (2000).
35. Meyer, C. P. & Paulay, G. DNA Barcoding: Error Rates Based on Comprehensive Sampling. *PLOS Biol.* **3**, e422 (2005).
36. Wickham, H. ggplot2: Elegant Graphics for Data Analysis. Springer-Verlag (2016).
